# Coupling of tactile LFP signals between mouse cortex and olfactory bulb

**DOI:** 10.1101/144279

**Authors:** Ana Parabucki, Ilan Lampl

**Author notes:** Correspondence to: Ilan Lampl.

## Abstract

Local field potentials are an important measure of brain activity and have been used to address various mechanistic and behavioral questions. We revealed a prominent whisker evoked local field potential signal in the olfactory bulb and investigated its physiology. This signal, dependent on barrel cortex activation and highly correlated with its local activity, represented a pure volume conductance signal that was sourced back to the activity in the ventro-lateral orbitofrontal cortex, located a few millimeters away. Thus, we suggest that special care should be taken when acquiring and interpreting LFP data.

## Introduction

Broad-band field potentials (FPs) are important and powerful tools when investigating and interpreting brain function. The reasons to believe so are ranging from the evidence of coupling/correlated local neuronal activity with FPs (Haider et al., 2016; Manning et al., 2009) to ability to play against the computer or overcome physical impairments through brain machine interfaces (Lebedev and Nicolelis, 2006). Thus, it is not a surprise that, in the last decade the recording of local field potentials (LFP) (Kajikawa and Schroeder, 2011) has gained increasing popularity in basic and applied electrophysiology. However, there is a uncertainty over LFP regarding its locality (Katzner et al., 2009; Kreiman et al., 2006; Schroeder et al., 1992) and its possible contamination by volume conductance coming from the distant generators (Buzsáki et al., 2012; Herreras, 2016; Kajikawa and Schroeder, 2011; Whitmore and Lin, 2016).

Here we observed whisker-dependent field potential in the olfactory bulb (OB), obtained using standard LFP recording techniques, and provide definite evidence that this signal is entirely a product of the volume conductance. Therefore, our findings add on to the rising concern regarding interpretation of LFP data, and consequently urge for careful experimental approach when dealing with LFPs (Herreras, 2016).

## Results

### Whisker stimulation evokes LFP response in mouse olfactory bulb

Using a glass micropipette, we recorded LFP from the OB of anesthetized mice while stimulating whiskers (usually 2-3), ipsi-(n=8, 4 mice) or contralaterally (n=12, 5 mice), with respect to the recording site (Figure 1A-B). Independent of the side, whisker stimulation caused a prominent and rapid LFP response in the OB. No differences were observed between ipsi- and contralateral responses in terms of amplitude (0.12 ± 0.02 vs 0.11 ± 0.02 mV, p = 0.97, two-sided Mann-Whitney U-test), onset latency (44 ± 6.31 vs 41.08 ± 4.01 ms, p = 0.88, two-sided Mann-Whitney U-test), and their respective distributions (Kolmogorov-Smirnov test). Therefore, only data from contralateral stimulation will be presented further. Neuronal response in the somatosensory system adapts in response to repeated tactile stimulation. Therefore, we examined the frequency response of the LFP signal (Figure S1), by stimulating whiskers at different frequencies. Observed responses adapted rapidly when whiskers were stimulated at 10 Hz, much less at 3 Hz and almost none at 1 Hz. Since cells in the somatosensory cortex also adapt rapidly (Katz et al., 2006) we next investigated if the response observed in the OB is correlated trial by trial with the response in the barrel cortex (Figure 1C). Simultaneous LFP recordings in the BC (layer 4 and 5) and OB (n = 12, 4 mice) revealed that OB whisker-evoked LFP response was delayed by 23.35 ± 1.19 relative to BC response, and strongly correlated to it in both onset (r^2^ = 0.58, p = 0.004) and amplitude (r^2^ = 0.87, p = 10^−5^). LFP signals in the OB were evoked only due to whisker stimulation. Air puffs to the back and an auditory stimulus (galvanometer click) did not elicit any response (Figure 2B). Since changes in respiration can affect neuronal responses in the OB (Carey and Wachowiak, 2011), we asked whether the observed LFP responses to whisker stimulation in the OB depend on the sniff cycle. Therefore, we recorded LFP simultaneously with the breathing cycle in three mice (Figures 2B-C, Methods) and observed no changes in the sniff cycle or its phases due to whisker stimulation (whole sniff cycle 401.85 ± 144.15 ms, p = 0.82, inhalation 103.9 ± 14.8 ms, p = 0.69 and exhalation 297.97 ± 135.05 ms, p = 0.83 One-way ANOVA).

**Figure 1.**
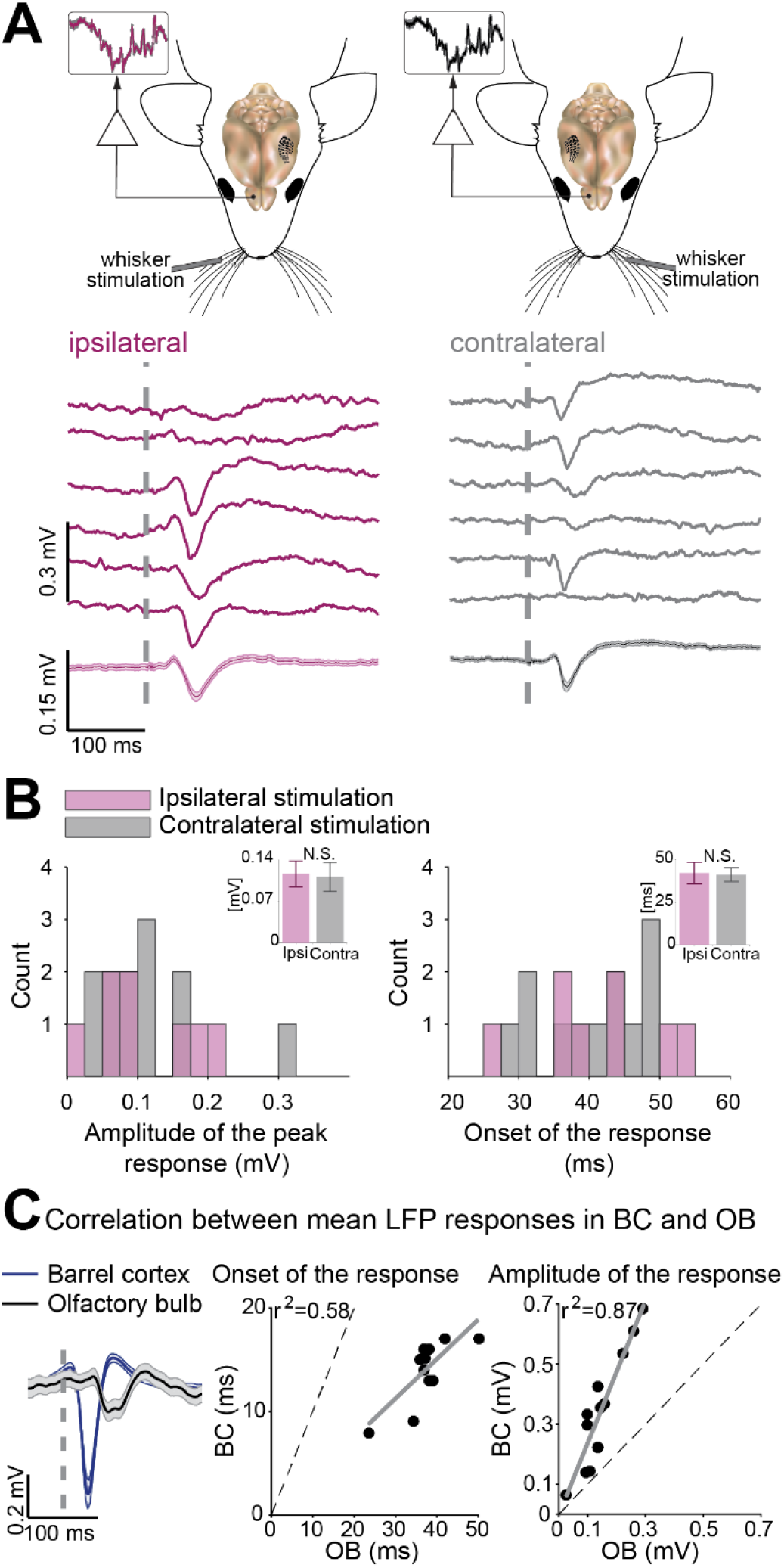
Whisker stimulation evokes LFP response in mouse olfactory bulb. (**A**) Representative ipsilateral (pink traces) and contralateral (dark-gray traces) LFP recordings in the OB with respect to the stimulated whisker (depictured in the in the mouse models above). The upper traces represent exemplary traces, while the lowest one is the mean with SEM given as shaded area. (**B**) The distribution of amplitudes (left) and latencies (right) in the population. Each bar represents the number of mean responses (contralateral = 12, ipsilaterlaral = 8) that fall under the given amplitude or time range, respectively. Insets represent the comparison of ipsi- and contralateral means of all experiments, with error bars (SEM). (**C**) On the left are exemplary average traces of simultaneously recorded responses in BC and OB. Correlation between mean whisker-evoked LFP responses from BC and OB is determined for onset (left, r^2^=0.58, p=0.004) and amplitude (right, r^2^=0.87, p=10^−5^) of the response (n=12). Onset of WS is depictured with dashed line. SEM are given as error bars and shaded areas. See also Figure S1.

**Figure 2.**
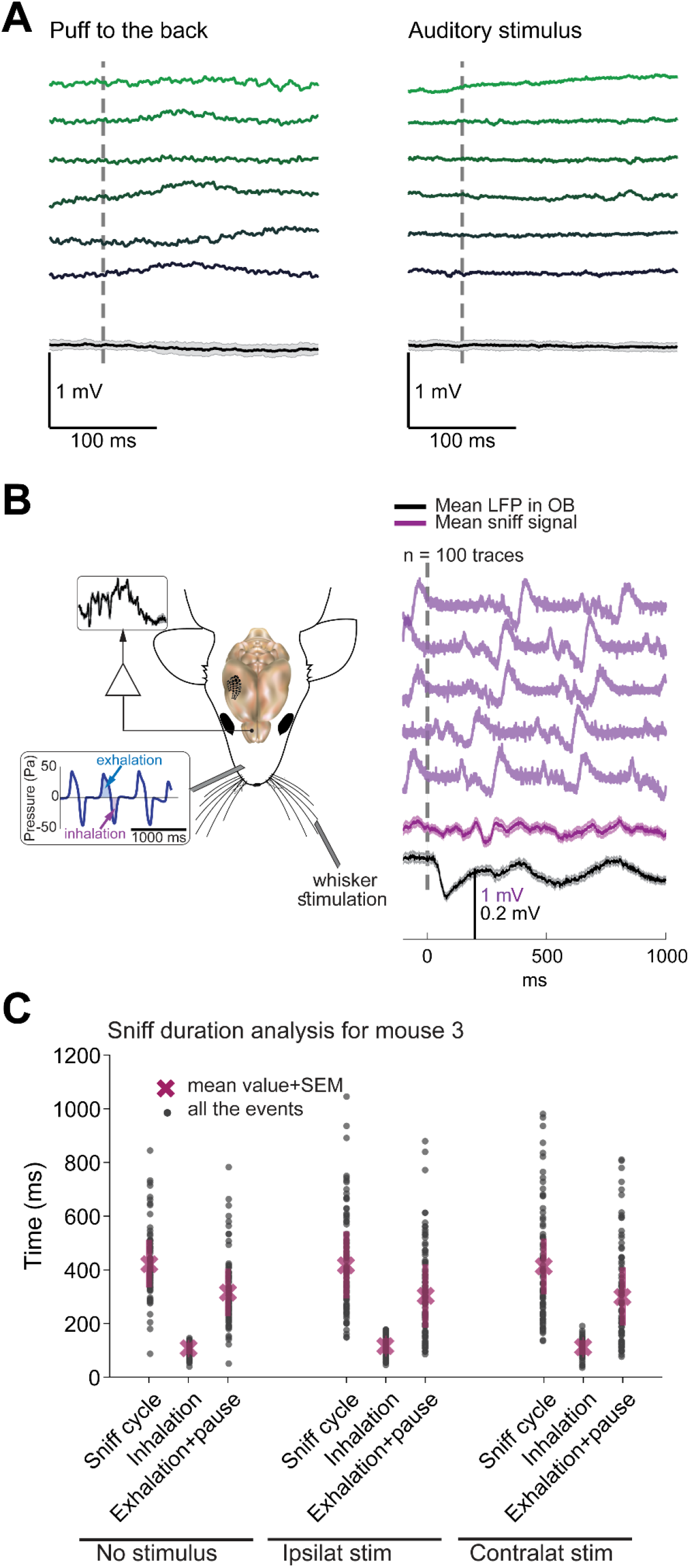
LFP sensory responses seen in the olfactory bulb are specific for whisker stimulation. (**A**) Left panel represents LFP signal recorded in the olfactory bulb, where mouse was presented with air-puffs to the back. On the right panel are depictured LFP signals from the OB while mouse was exposed to the auditory stimuli, matching those that emerge during the experiments (galvanometer click). Upper traces are exemplary, individual traces, while the bottom ones are the mean of the response. (**B**) Sniff analysis was performed on data obtained as depictured in diagram on the left. On the right panel, upper traces represent individual traces of the sniff signal in anesthetized mouse (light purple), while the average sniff signal is given below (purple trace). The black trace represents the mean LFP recorded from OB in the same experiment. (**C**) Duration of the complete sniff cycle and its phases (inhalation and exhalation + pause) is presented as mean value + SEM, and black dots represent individual values. Comparison was carried among fourth sniff in control (no stimulation), ipsi- and contralateral whisker stimulations experimental paradigms (for details, see Methods). Onset of the stimulus is depictured with dashed line. SEM are given as error bars and shaded areas.

### Whisker evoked-LFP in olfactory bulb depends on the BC activity

To test whether the LFP response observed in the OB upon whisker stimulation depends on the BC, we employed optogenetic and pharmacological tools (Figure 3 and Figure S2). First, we used Thy1-ChR2 mice which express ChR2 in layer 5 neurons and illuminated the BC with blue light. This induced a strong LFP response in the OB, which strongly adapted using repetitive light stimulation at 8 Hz. In addition, substituting the last light stimulation with whisker stimulation led to significant attenuation of whisker evoked LFP response in the OB, indicating that the adaptation depends on at the common origin – the barrel cortex (Figure 3A, T7 = 0.04 ± 0.01, G =, 0.11 ± 0.01, n = 4, p=0.02, Wilcoxon signed-rank test). We confirmed these observations using electrical stimulation of layers 4/5 of the BC (Figure S2A).

**Figure 3.**
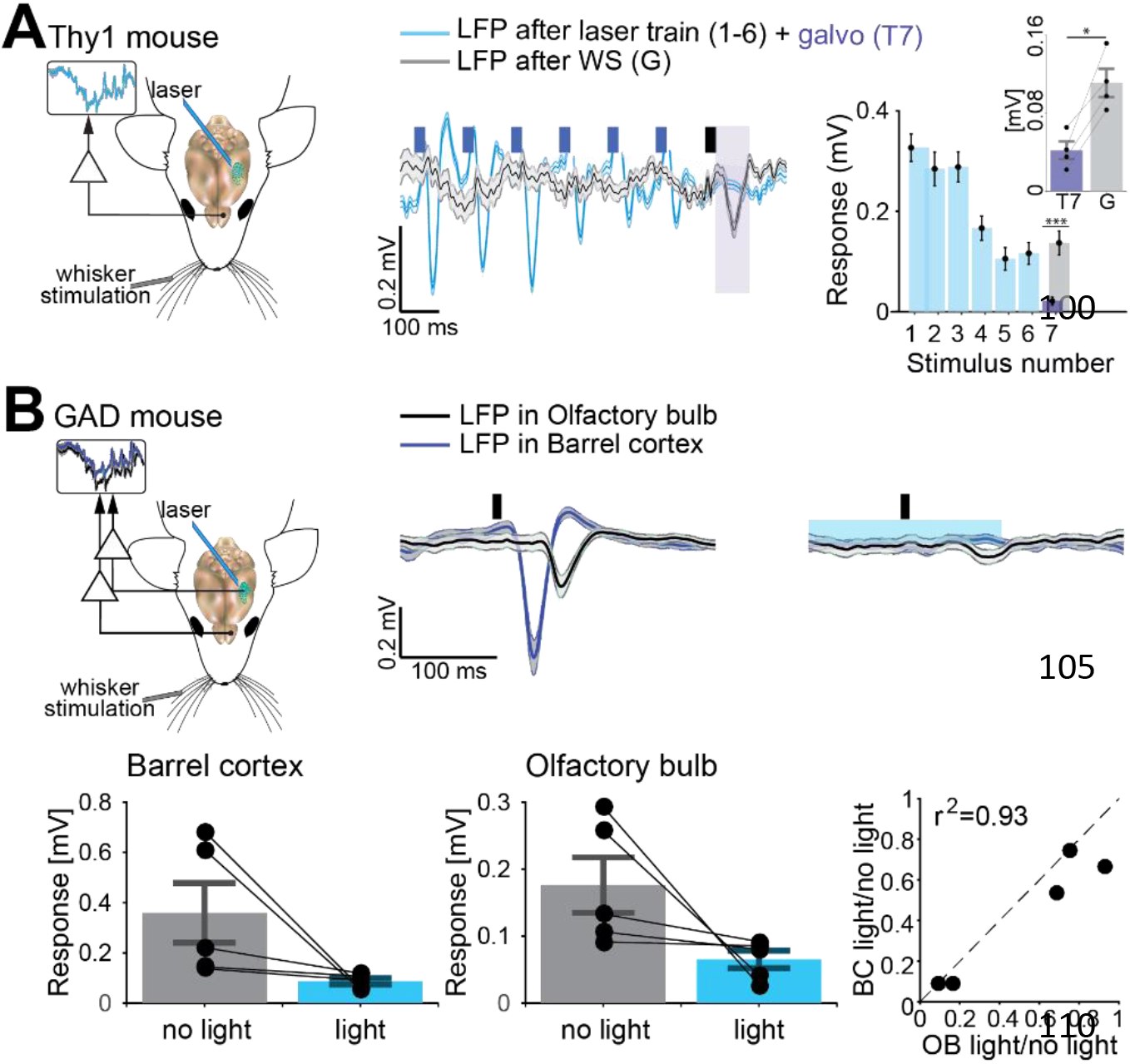
Whisker-evoked LFP signal in OB depends on barrel cortex. (**A**) Light activation of the barrel cortex in Thy1-ChR2 mice induced responses in the olfactory bulb (mouse diagram). Responses to six consecutive light stimulations of the BC (t_stim = 10 ms, freq = 8 Hz, blue traces and bars) followed by a galvo stimulation of the contralateral whisker (purple traces and bars) were compared to galvo whisker stimulation without optogenetic pre-pulses (grey traces and bars (T7 = 0.02 ± 0.01 mV, G = 0.14 ± 0.02, p = 0.0005). Inset represents the population statistics, where responses to whisker stimulation with (T7) and without (G) preceding light trains were compared (T7 = 0.04 +− 0.01, G = 0.11 ± 0.01, n = 4, p=0.02). See also Figure S2A. (**B**) Paired LFP recordings in the BC (dark-blue traces) and OB (black traces) in GAD-ChR2 mice (mouse diagram) showed attenuated whisker-evoked LFP responses after light inhibition of the BC, in both structures. Below: population analysis of response amplitude to the light in BC (before light 0.36 ± 0.12 mV, and after light 0.09 ± 0.01 mV, p = 0.008) and OB (before light 0.18 ± 0.04 mV, and after light 0.07 ± 0.01 mV, p = 0.02), (left and middle panel, respectively), presented as the mean amplitude of the whisker-evoked LFP response per each recording pair (black bars no light, blue bars during light). Right panel shows change in BC versus OB, after light-inhibiting the BC SEM are given as error bars and shaded areas. See also Figure S2B.

Next, we used GAD-Cre mice which express ChR2 in GABA neurons to silence the BC (Malina et al., 2016) while recording simultaneously from both BC and OB. Whisker stimulation induced responses in both structures, while light inactivation of the barrel cortex attenuated those responses in both BC (before light 0.36 ± 0.12 mV, and after light 0.09 ± 0.01 mV, p = 0.008) and OB (before light 0.18 ± 0.04 mV, and after light 0.07 ± 0.01 mV, p = 0.02, Wilcoxon signed-rank test) where the percentage of attenuation was highly correlated (r^2^ = 0.93, p = 0.0083, 5 mice, Figure 3B). These results were in accordance with observation upon topical application of Muscimol (40 μL, 2.5 mM) to the BC, which significantly reduced amplitude of the whisker-evoked LFP response in the OB (Figure S2B).

Interestingly, intracellular recording from 15 neurons in the OB (Figure S3), aimed to find the cellular correlates of the whisker-evoked LFP responses, did not show any response to whisker stimulation. This clearly called for further investigation of the origin of the whisker evoked OB LFP response.

### Whisker-evoked LFP in olfactory bulb is passively conducted from ventrolateral orbital cortex

In order to trace the origins of the whisker-evoked LFP in OB, we recorded from different brain structures (Figure S4). Surprisingly, paired recordings in the ventro-lateral orbitofrontal cortex (vlOrb) and OB showed no significant difference in the onset of the whisker-evoked LFP between these two structures (1.31 ± 0.35 ms, n = 9, 3 mice, Figure 4A). Next, we performed multichannel recording of the LFPs in the OB followed by inverse current source density (iCSD) analysis (Figure 4B, Figure S5, Methods). LFP traces recorded in different channels did not exhibit any difference in time lag or amplitude with the depth of the recording point. Accordingly, iCSD analysis could not identify sinks and sources of the current (unlike in the BC, see Figure S5B), strongly suggesting that the observed signal does not reflect synaptic transmission within the OB. In contrast, iCSD analysis of recordings from the vlOrb show clear sink and source (Fig. 4C). Moreover, the whisker evoked LFP responses in the OB depended on the response in the vlOrb, as confirmed by injection of ~10 – 15 μL Muscimol (2 mM) bilaterally to the vlOrb (Figure S5C-D). Approximately 20 minutes following the injection, the whisker evoked LFP response in the OB was completely attenuated. Together, these recordings strongly suggest that whisker-evoked OB LFP response reflects a volume conductance signal arising from the vlOrb.

**Figure 4.**
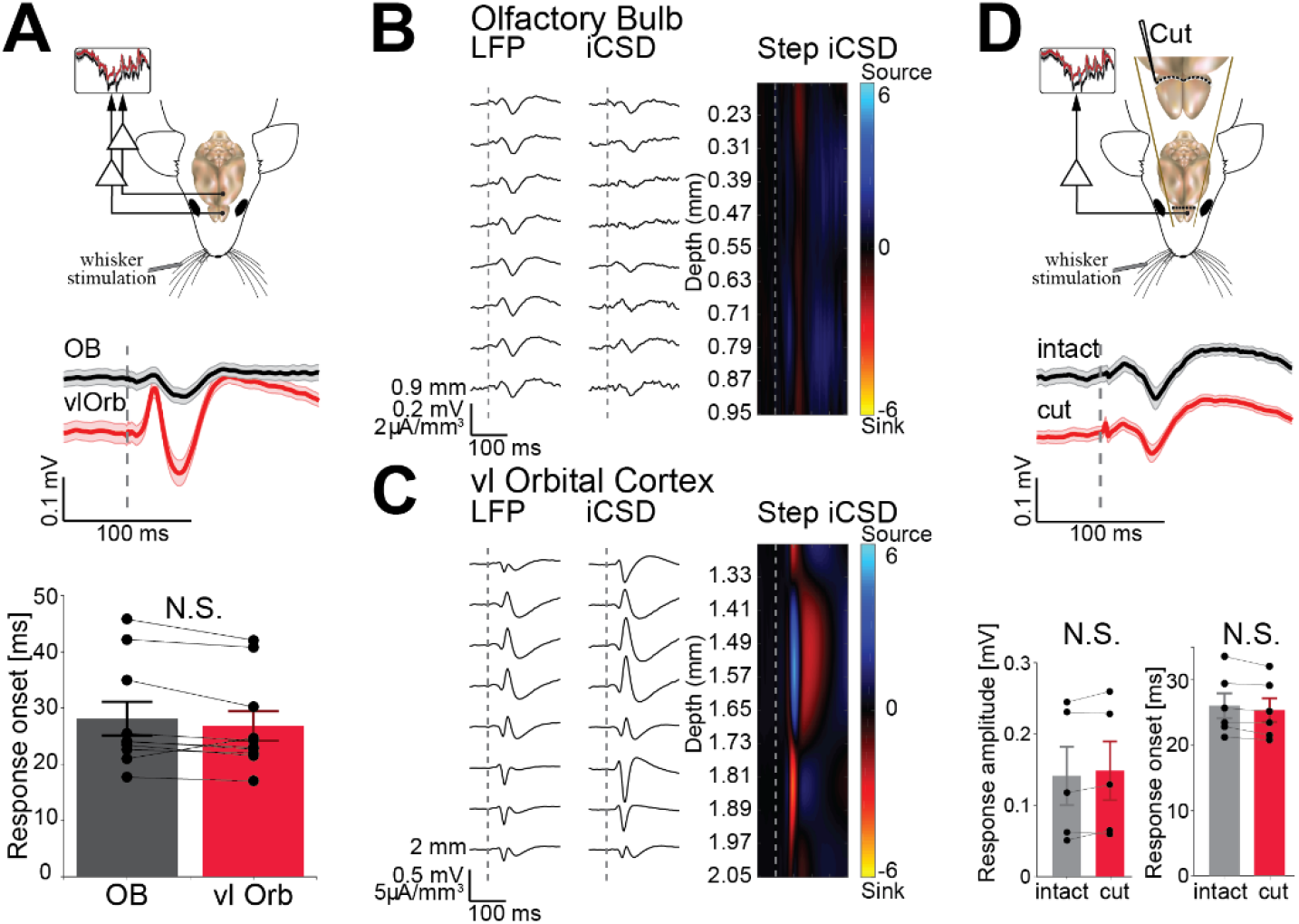
Whisker-evoked LFP response in OB spreads from cortex as a volume conductance signal. (**A**) The diagram represents the experimental setup for double LFP recordings from OB (black traces and bars) and vlOrb (red traces and bars), while below is given mean LFP response from the representative experiment. Response onset in OB and vlOrb is presented in form of bar plot (n=9) in the bottom panel. (**C-D**) Middle panels represent mean whisker-evoked LFP responses recorded with a multichannel silicone probe in OB and vlOrb, respectively. The middle panel depictures iCSD profiles calculated from the given LFPs, which are presented in a form of colormap on the right; negative deflections are colored red and positive in blue (current sinks and sources, respectively). (**D**) Recordings in the OB before and after disconnecting the OB are given in black and red traces, respectively. The diagram above expands the lower, illustrating the position of the cut. Exemplary mean responses are given in the middle panel, while bar plot depictures amplitude of the response in the population (n=5). See also Figures S4, S5 and S6. SEM are given as error bars and shaded areas.

Finally, our hypothesis was confirmed by recording whisker-evoked LFP (3 mice) following separation of the OB from the rest of the brain by a precise cut (Figure 4D). As expected for the case of volume conductance, cutting all the connections between OB and the rest of the brain did not attenuate the LFP response to the whisker stimulation; both the the amplitude (before 0.14 ± 0.04 mV and after the cut 0.15 ± 0.04 mV, p = 0.84, Wilcoxon signed-rank test) and the latency of the signal onset (before 25.98 ± 1.89 ms, and after the cut 25.3 ± 1.81 ms, p = 0.73, Wilcoxon signed-rank test) remained unaltered.

Since ground placement can be a major issue when recording extracellular signals(Kajikawa and Schroeder, 2011; Whitmore and Lin, 2016), we tested different positions of the grounding wire which are classically used in the field (Figure S6). The whisker-evoked LFP in the OB persisted in the various recording configurations and disappeared only when the ground electrode was placed within the OB, in the vicinity (< 1 mm) of the recording electrode. The above results indicate that the whisker-evoked LFP recorded in the OB is mediated via volume-conductance spread of the LFP signal in the vlOrb, propagating rostrally over 2-4 mm to the OB.

## Discussion

There is a growing interest in studying cortical and subcortical regions using LFP recordings, suggesting that volume conductance effects should be further investigated in detail. Our results, demonstrating a pure volume conductance tactile signal in the OB that originates in cortex, make the OB is a good candidate due to its position relative to the rest of the brain and the possibility to functionally disconnect it while the animal remains alive (Ito et al., 2014). Additionally, our understanding of the LFP signal in the brain is especially important for studies in awake animals and humans, where we may expect the contamination due to volume conductance to be even more pronounced. In fact, in their recent paper Marmor and colleagues (Marmor et al., 2017) show that LFP recording in human subthalamic nucleus using common monopolar electrode contributes to significant contamination of the signal. Authors also suggest that this volume conductance might spread from cortex, located more than 80 mm away.

In summary, we demonstrated that a pure volume conductance spread of LFP signal can be detected millimeters away from its origin. Our study suggests therefore that interpretation of LFP signals alone might lead to erroneous conclusions pertaining to the functional connectivity of different brain regions. Although extremely helpful, LFPs should not be recorded from single channels without further confirmation. This should be based on single unit activity and appropriate multichannel probes that enable CSD analysis (Pettersen et al., 2006) and profound mathematical approaches (Einevoll et al., 2013; Lindén et al., 2011) for complete and reliable interpretation of the observed LFP signal.

## AUTHOR CONTRIBUTIONS

A.P. designed and performed the experiments. I.L. designed and supervised the project. A.P. and I.L. wrote the manuscript.

## Acknowledgments

This work was supported by DFG-SFB 1089. We thank all the members of the I.L. laboratory and especially Yonatan Katz, Katayun Cohen-Kashi Malina and Michael Sokoletsky for their helpful contributions. We especially thank Gilad Silberberg for his helpful comments on the full manuscript.

## Material and Methods

### Animals

All the procedures involving animals were reviewed and approved by the Weizmann Institute Animals Care Committee. In experiments, we used 8–14 weeks old mice of either sex housed up to five in a cage with a 12/12 h dark/light cycle. The following mouse strains were used: C57BL/6 mice, GAD-CRE mice (JAX #010802) crossed with a ChR2 reporter strain (JAX #012569), and Thy1-ChR2-YFP (JAX # 007612).

### Animal preparation

Mice were dosed with a mixture of ketamine (90 mg kg^−1^, Ketamidor, Richter Pharma AG, Wels, UK) and domitor (2 mg kg^−1^, Domitor, Vetoquinol UK Limited, Buckingham, UK) intraperitoneally. The skin was removed and the skull exposed and cleared, followed by mounting a metal head-plate over unused hemisphere using dental cement (Lang dental, Wheeling, IL, USA). A craniotomy (~1-2 mm in diameter) was made above the olfactory bulb (various positions, ranging frombregma: +6 until +4, lateral: +0.25), barrel cortex (bregma: −0.70, lateral: 3.30) and/or orbitofrontal cortex (bregma: +2.6, lateral: 1.00, depth: 0.8-2.10) and a portion of the dura mater was carefully removed. The craniotomy was kept moisturized with artificial cerebrospinal fluid (ACSF) containing (in mM): 124 NaCl, 26 NaHCO3, 10 glucose, 3 KCl, 1.24 KH2PO4, 1.3 MgSO4 and 2.4 CaCl2. The heart rate (250–450 beats per minute) was monitored throughout the experiments and body temperature was kept at 37 °C using a heating blanket and a rectal thermometer.

For the experiments in which the bulb was cut (3 mice), a full bilateral craniotomy was carefully made to expose the entire caudal part of the olfactory bulb. After that, the control recording was performed, the position of the recording noted and the pipette was retracted. Olfactory bulb tissue was then cut and completely separated from the rest of the brain using a surgical scalpel. The incision was washed from blood and debris with ACSF and any bleeding was stopped using hemostatic gelatin sponge (Cutanplast, Milano, Italy). After that, the recording pipette was lowered to the approximate position of the control one, and the LFP was recorded.

### Whisker stimulation, optogenetic, electrical and pharmacological manipulations

Whiskers were trimmed to a length of 10–20 mm. Whiskers were inserted into 21G needle attached to a galvanometer servo motor with a matching servo driver and controller (6210H, MicroMax 677xx, Cambridge Technology Inc., Cambridge, MA, USA). A fast-rising voltage command was used to evoke a fast whisker deflection with a constant rise time of ~1 ms followed by a 20 ms ramp-down signal. The stimulation velocity and the corresponding deflection amplitude (~50 mm s^−1^, 145 μm amplitude) were used to evoke LFP responses in given brain structures.

To activate ChR2, LED light source of 460 nm (Prizmatix Opt-LED-460, Givat-Shmuel, Israel) was coupled to a bare optical fiber (200 μm diameter, 0.22NA; ThorLabs M25L05, Newton, NJ, United States) and placed above the cortex. The LED was driven by an analogue output from our acquisition system (National Instruments, Austin, TX, USA) for either 10 ms in case of pulse activation in Thy1-ChR2 mice or 200 ms in case of silencing the barrel cortex in GAD-Cre mice (starting 100 ms before whisker stimulation). The intensity of the light was around 7 mW at the tip of the fiber.

For electrical stimulation, we stereotaxically placed a concentric needle electrode (30G; Alpine Biomed ApS, Skovlunde, Denmark) in the barrel cortex at a depth of 350-700 μm. Either single or multiple (20 Hz) current pulses (amplitude of 400 μA, duration of 0.5 ms) were injected by an isolation unit (ISO-Flex; A.M.P.I. Instruments, Jerusalem, Israel).

To silence the barrel cortex and ventro-lateral orbital cortex pharmacologically, we used muscimol hydrobromide (Sigma-Aldrich, St. Louis, MO, USA) – a potent agonist of GABA-A receptors. In the case of silencing the barrel cortex (2 mice), muscimol was applied topically (40 μL, 2.5 mM) into a previously made well to prevent dissipation of the liquid and drying of the tissue. When muscimol was used to silence vlOrb cortex (2 mice), a glass micropipette with an ACSF solution containing muscimol (~15 μL, 2.5 mM; Sigma-Aldrich, St. Louis, MO, USA was inserted through a cranial window at the coordinates designated above, at a depth of ~ 1.2 mm from the cortical surface. The solution was injected by hand at ~100 mbar (monitored with manometer, Lutron, Coopersburg, PA, USA) for 1 min through a pipette with tip diameter ~30 μm. After each injection and the removal of the pipette from the brain, pressure was reapplied to confirm that the pipette tip was not clogged.

Whenever applicable, trials were delivered pseudo-randomly. To avoid synchronization with sniffing cycle in anesthetized mice, inter-trial intervals were pseudo-randomly and trial-to-trial altered in the range of 2-4 s. Each condition included in the analysis of this study was repeated no less than 30 times.

### Electrophysiology and sniff recordings

Borosilicate micropipettes were pulled to produce electrodes with a resistance of 4–10 MΩ when filled with an intracellular solution containing the following (in mM): 136 K-gluconate, 10 KCl, 5 NaCl, 10 HEPES, 1 MgATP, 0.3 NaGTP, and 10 phosphocreatine (310 mOsm/L). Grounding wire was positioned in the mouse neck area (unless stated differently), enabling contact with the skull and surrounding tissue. However, the same results were obtained when the animal was grounded at different locations (Figure S6, ground position 1, 3 and 4, as well as grounding during multichannel recordings described below). Single electrode field potentials and intracellular signals were acquired using a Multiclamp-700B or Axoclamp-900B amplifier (Molecular Devices, Sunnyvale, CA, USA) and band passed at 4 kHz before being digitized at 10 kHz. Cell potentials were obtained in current-clamp mode. A custom-built interface written in LabVIEW (National Instruments, Austin, TX, USA) was used for data acquisition.

NeuroNexus silicone probes (A1x16-5mm-150-703, NeuroNexus, Ann Arbor, MI, USA) were used for obtaining LFP profiles, and consequently inverse current source density (iCSD) analysis. Data was obtained from 8 neighboring channels, 100 μm apart and with a measured resistance of ~1.5 MΩ, recorded using LabVIEW and Intan acquisition systems (RHD2000, Intan Technologies, Intan Technologies, Los Angeles, CA, USA) at 20 KHz sampling frequency. In addition, a small screw was implanted in the skull above the V1 area and connected to the grounding wire, while a reference wire was implanted in the cerebellum.

Recording depth in olfactory bulb ranged between 100 and 1000 μm, and in some cases cells were obtained from deeper regions. Recordings in the barrel cortex were performed in layers 4 and 5 (whereas recordings within 250-500 μm range were classified as layer 4, while those within 500-700 μM were classified as layer 5).

In mice where the breathing cycle was monitored, a small hole was drilled into the nasal bone and a ~7 mm long stainless-steel cannula (gauge 23) was inserted and fixed using dental cement. The cannula was then connected to a pressure sensor with polyethylene tubing (A-M systems, Sequim, WA, USA). Sniffing caused changes in pressure which were detected by pressure sensor (Honeywell, Morris Plains, NJ, USA) and a homemade preamplifier circuit. The output analog signal was recorded in parallel with the electrophysiological data.

### Data analysis

The LFP were separated from the high frequency signal by bandpass filtering at 0.1 to 100 Hz, while the intracellular recordings were low-passed at 4 kHz before being digitized at 10 kHz. All recordings were analyzed using custom programs written in MATLAB (MathWorks, Natick, MA, USA). We smoothed the raw traces using a symmetric Savitzky–Golay filter with a first-order polynomial and a window size of 21 points. The amplitude of the LFP or cell response was measured as the difference between the peak of response over 100 ms after the stimulation and the mean baseline value obtained over 10 ms before stimulation.

Sniffing traces were down-sampled to 1 kHz and filtered in the range of 0.1-30 Hz. The phases of the sniff were acquired using a similar approach to one described by Shusterman (Shusterman et al., 2011) In brief, the onset and offset of the inhalation were detected by determining zero crossings of parabola fit. Inhalation onset was defined as the first, and inhalation offset as second zero crossing of the parabola. The interval between the previous inhalation offset and next inhalation onset was defined as exhalation + pause.

Coefficients of correlation were calculated using an integrated Matlab function.

The inverse CSD (iCSD) method is assumed to have cylindrical symmetry and localized in infinitely thin discs. It is based on the inversion of forward electrostatic solutions and to calculate it we used a slightly modified version of CSDplotter toolbox (Pettersen et al., 2006). The same toolbox was used when creating step iCSD (colormaps), which assumes step-wise constant CSD between the electrode contacts. In all the recordings, the estimates at the top and bottom electrode channels were provided by the method of Vaknin (Vaknin et al., 1988) and the default filter used was the Hamming filter (center weight: 0.54, neighboring weight: 0.23). Assumed tissue conductivity was 0.3 S/m, same as the conductivity above the tissue (ACSF). Activity diameter was assumed to be 0.5 mm in all experiments.

### Statistics

All values are indicated as mean ± s.e.m. Significance was determined at a significance level of 0.05 in all tests. The Kolmogorov-Smirnov test was used to check whether the cumulative distribution of responses amplitude and onset were significantly different among ipsi- and contralateral groups (Figure 1B). For comparisons in which the number of samples in the two conditions was not equal, we used the two-sided Mann-Whitney U-test (Figure 1B inset), whereas for paired data we used the two-sided Wilcoxon signed-rank test (Figure 3A-B, Figure 4A-D, Figure S2B). In all cases, automated analysis was performed.

### Data availability

The data that support the findings of this study are available from the corresponding author on request.

